# CRISPR/Cas9-mediated editing of the *ADE2* locus in *Saccharomyces cerevisiae*: targeted mutagenesis and homology-directed repair

**DOI:** 10.1101/2025.03.30.646154

**Authors:** Essa Jarra

## Abstract

The CRISPR/Cas9 system, derived from the adaptive immune defence of bacteria and archaea, has emerged as a powerful tool for genome engineering by enabling precise, site-specific modifications. Type II bacterial CRISPR systems recognize and cleave DNA in various microorganisms using the RNA-guided Cas9 endonuclease. We present a streamlined and efficient approach for genome editing in two strains of *Saccharomyces cerevisiae*, focused on targeted mutagenesis of the *ADE*2 gene of the BY4741 strain and homology-directed repair of the *ade2* locus of the W303-1A strain, involving a single-step transformation strategy utilizing the pML104 plasmid, which encodes the Cas9 endonuclease and a user-defined single-guide RNA (gRNA) complementary to the target site. In the BY4741 (wildtype) strain, pML104-gRNA components achieved successful *ADE2* disruption rates of 40% (95% CI: 0.10–0.70), confirmed by adenine auxotrophy screening and Sanger sequencing. Additionally, co-transformation of gRNA plasmid and donor repair templates in the W303-1A (*ade2* mutant) strain also resulted in 40% recombination efficiency, restoring the function of the mutant gene to allow activation of *de novo* purine biosynthesis activity when cultured on SC-Ade medium. This optimized, cost-effective CRISPR/Cas9 method enhances editing efficiency in *S. cerevisiae*, providing a robust and practical platform for high-precision genetic modifications with broad applications in molecular biology and biotechnology.

## Introduction

The Clustered Regularly Interspaced Short Palindromic Repeats (CRISPR) and CRISPR-associated (Cas) system constitute an adaptive immune defense mechanism in bacteria and archaea, protecting against invading bacteriophages and plasmids [1, 2]. Six distinct types of CRISPR-Cas systems [Types I-VI] [3] have been identified in diverse bacterial species. Among these, types I–III are the most extensively studied [4, 5], while types IV–VI have been more recently characterized [6]. The Type II CRISPR system from *Streptococcus pyogenes* is particularly notable for its ability to recognize and cleave DNA in various microorganisms via an RNA-guided Cas9 endonuclease [7]. Each type of CRISPR/Cas system consists of multiple *Cas* genes located adjacent to an accompanying CRISPR array of individual repetitive sequences (repeats) interspaced with short unique sequences (spacers). The spacers are segments of genetic elements derived from previous bacteriophages and plasmids (protospacer) infections [1, 8]. Upon transcription, the CRISPR locus is processed into mature CRISPR RNAs (crRNAs), each carrying an individual spacer sequence [8]. The crRNA complexes with a trans-activating crRNA (tracrRNA) to form a functional guide that directs Cas9 to cleave target DNA sequences carrying a complementary protospacer sequence [8]. To facilitate genome editing applications, Jinek et al. [4] designed a chimeric guide RNA (gRNA) that combines the features of and mimics the crRNA-tracrRNA complex, streamlining CRISPR-based genome engineering.

The sequence-specific cleavage activity of Cas9 requires the presence of a protospacer adjacent motif (PAM) with the sequence 5’-NGG-3’ immediately downstream of the target DNA sequence [2]. Upon recognition, Cas9 introduces a double-strand break (DSB) through the activity of its RuvC and HNH nuclease domains, which respectively cleave the non-complementary and complementary DNA strands relative to the gRNA [2, 4]. The resulting DSBs can be repaired by the error-prone non-homologous end joining (NHEJ) pathway, often introducing genomic mutations that lead to gene disruption [9]. Alternatively, in cells favoring homologous recombination, precise genome editing can be achieved when Cas9 is co-introduced with a donor DNA repair template, facilitating homology-directed repair (HDR) at the break site [10].

CRISPR/Cas9 has emerged as an exceptionally versatile genome engineering tool, allowing precise genetic modifications across a wide range of organisms [5]. Compared to other genome-editing technologies such as zinc-finger nucleases (ZFNs) and transcription activator-like effector nucleases (TALENs), the CRISPR/Cas9 system offers superior specificity, as target recognition is solely determined by base-pair complementarity with the easy-to-design gRNA, making it more accessible and efficient for genetic manipulation [11]. Consequently, CRISPR has been widely adopted for applications in molecular biology, medicine, and biotechnology. Notably, CRISPR-based tools have been employed for targeted gene modification in human cells [12], generation of animal models for studying human genetic diseases [13-15], and genome-wide knockout screening in human cells [16]. Additionally, others have engineered Cas9 protein to function without its nuclease activity for applications such as gene expression modulation [17], and as a gene-imaging tool [18]. More recently, a catalytically inactive Cas9 (dCas9) variant has been used in the presence of PAM-presenting oligonucleotides for programmable tracking of RNA in live cells without requiring genetic tagging [19].

*Saccharomyces cerevisiae* (budding yeast) is a well-established model organism for genetic studies, with approximately 31% of its genes having homologs in mammals [20]. Research utilizing yeast homologs of human genes provides useful insights into gene function, significantly improving our understanding of the genetic basis of human diseases. For example, studies of yeast homologs have contributed to the understanding of hereditary conditions such as nonpolyposis colorectal cancer (*MSH2* and *MLH1* in yeast) and Werner’s syndrome (*SGS1* in yeast) [21]. Additionally, the ability to match sequenced human disease genes to their homologs with known functions in *S. cerevisiae* has facilitated many functional studies [21]. The *ADE2* gene in *S. cerevisiae* encodes an enzyme involved in the conversion of phosphoribosylaminoimidazole (AIR) to phosphoribosyl-carboxy-aminoimidazole, catalyzing the sixth step of the *de novo* purine biosynthesis pathway [22]. The transcription of *ADE2* is tightly regulated by adenine availability, with gene expression being upregulated in the absence of adenine [23]. Mutations in *ADE2* disrupt this pathway, leading to the accumulation of AIR, a red pigment, which results in the formation of red-coloured colonies when grown on adenine-deficient media [24]. W303-1A and its derivative strains of *S. cerevisiae* carry adenine auxotrophic markers (*ade1* and *ade2*) and are widely used for genetic selection and screening applications [22].

In this study, we separately transformed a plasmid (pML104) encoding the Cas9 endonuclease into a wildtype (BY4741) and mutant (W303-1A) *S. cerevisiae* strains to assess its genome editing efficiency. A gRNA was designed to target the *ADE2* locus at nucleotide position 141, directing Cas9 to introduce a DSB at nucleotide position 146. In the wild-type strain, the DSBs were repaired via the non-homologous end joining (NHEJ) pathway, generating nonsense mutations in *ADE2*, which were phenotypically identified. Sequence alignment of PCR-amplified mutated alleles confirmed point mutations at the Cas9 cleavage site in BY4741 cells, verifying the effectiveness of CRISPR-mediated mutagenesis. Additionally, homology-directed repair (HDR) was demonstrated by co-transforming pML104-gRNA with single- and double-stranded donor DNA templates into the W303-1A mutant strain, successfully restoring the *ade2* locus to its wild-type sequence and introducing a mutation in the PAM to prevent further Cas9 activity. This study highlights the utility of CRISPR/Cas9 for precise and efficient genome editing in *S. cerevisiae* and provides a streamlined approach for site-specific mutagenesis and HDR in yeast.

## Materials and Methods

### Construction of pML104 and cloning of the gRNA

The construction and use of the pML104 plasmid (Addgene plasmid #67638; https://www.addgene.org/67638/), which contains the Cas9 gene (Figure 1), were described by Laughery et al. [10]. A 50µL aliquot of overnight *Swa*I-digested pML104 (50ng/µL) was further digested with 2µL *Bcl*I (New England Biolabs) for 2 hours at 50ºC. The plasmid DNA was purified using the GeneJET PCR Purification Kit (Thermo Fisher Scientific) according to the manufacturer’s instructions.

**Figure 1.**
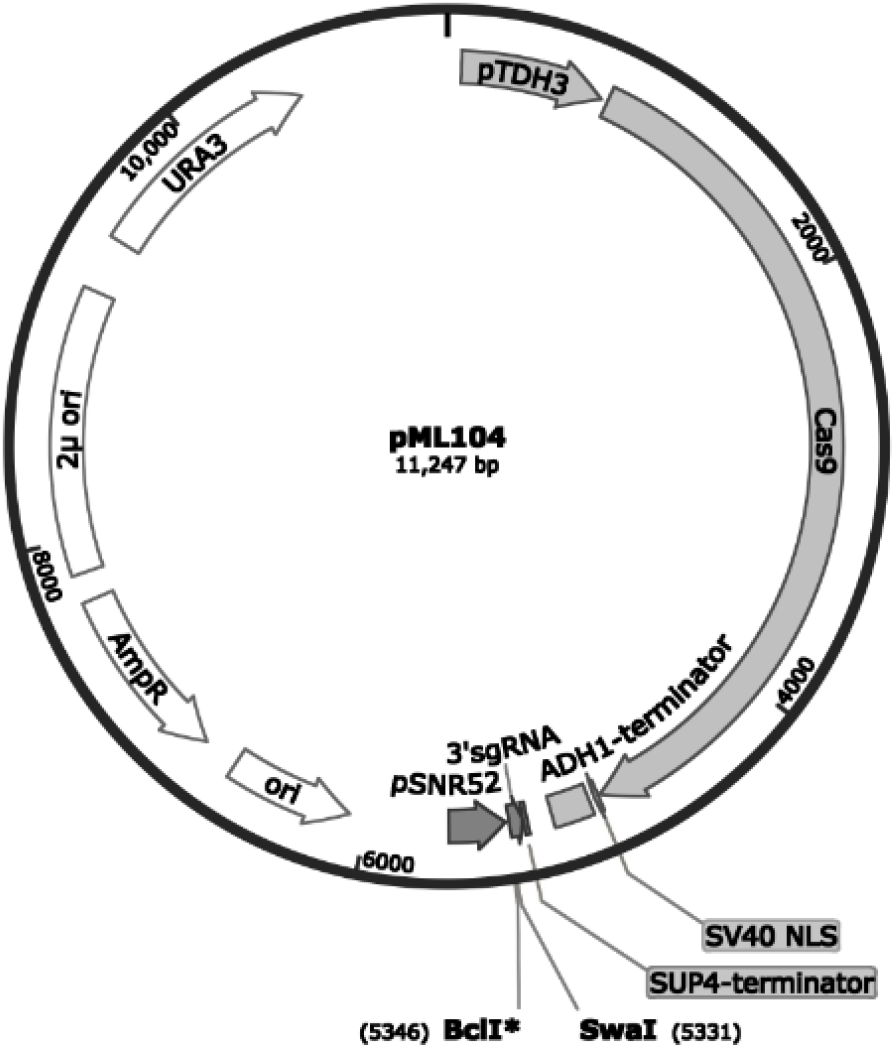
Schematic map of the pML104 plasmid, containing the Cas9 gene, the single-guide RNA (sgRNA) expression cassette, and a URA3 selectable marker flanked by restriction enzyme sites that facilitated gRNA cloning for one-step CRISPR/Cas9 editing in yeast. Retrieved from: https://www.addgene.org/67638/.

The gADE2_f and gADE2_r oligonucleotides (Eurofins Genomics) encoding the gRNA sequence for targeting the *ADE2* gene were designed using the Yeast gRNA Search Engine [10], and high-quality target sites were identified using Chopchop [25]. The gADE2_f and gADE2_r oligonucleotides were annealed by mixing 2µL of each with 10µL of 10x ligation buffer (Sigma-Aldrich, UK) and 86µL of sterile water. See Supplementary Material Table S1 for sequences of oligonucleotides. The mixture was incubated at >90ºC for 5 minutes in a metal heat block, allowed to cool to room temperature, and diluted with 150µL of sterile water.

The annealed oligonucleotides were ligated into the digested pML104 by mixing 1µL of diluted annealed oligonucleotides, 7µL of purified digested pML104, 1µL of 10x ligation buffer (Sigma-Aldrich, UK), and 1µL of T4 DNA ligase (Sigma-Aldrich, UK). The ligation reaction was incubated overnight at 18ºC.

### Bacterial transformation and PCR verification of pML104-gRNA clones

For routine plasmid propagation, 9µL of the pML104-gRNA ligation mixture and 1µL of unmodified control plasmid (pML104) were independently transformed into 100µL of competent *E. coli* Top10 cells (Supplementary Material Table S1). The cells were incubated at 42ºC for 60 seconds and then recovered by adding 1mL of enriched lysogeny broth (LB) (1% tryptone, 0.5% yeast extract, 1% NaCl in water). Cultures were incubated at 37ºC for an additional 30 minutes. The bacterial cells were pelleted by centrifugation at 3,000 rpm for 5 minutes, the supernatant was discarded, and the pellets were resuspended in a residual drop of medium. Cells were transferred onto separate pre-prepared LB/Ampicillin (100mg/L) agar plates (1% Tryptone, 0.5% yeast extract, 1% NaCl and 2% agar in water) and incubated at 37ºC for 24 hours.

Four independent *E. coli* TOP10 transformants were selected and separately inoculated into liquid LB medium. The bacterial cells were pelleted by centrifuging 2ml of each culture at 13,000rpm for 1 minute. The supernatants were decanted, and all remaining media was removed. Plasmid DNA was extracted using the GeneJET Plasmid Miniprep Kit (Thermo Fisher Scientific) following the manufacturer’s instructions. Analytical PCR verification of the pML104-gRNA was performed for the four DNA extracts using 25µL reactions containing 12.5µL GoTaq Green MasterMix (Promega), 1µL each of gADE2_f and g104_check primers (Eurofins Genomics), 1µL plasmid DNA, and 9.5µL sterile water. The PCR amplified the region including the gRNA cassette and up to150 nucleotides upstream of the gRNA sequence. See Supplement Materials for PCR amplification cycle. The products were resolved via electrophoresis on a 1% agarose gel (0.75g agarose in 75mL 1x TAE buffer with 7.5µL SYBR Safe stain [Sigma-Aldrich, UK]) and visualized under UV illumination.

For sequence confirmation, pML104-gRNA DNA was amplified using ADE2_up and ADE2_sta primers (Eurofins Genomics), with cycling conditions listed in Supplementary Material. The amplicons were sequenced using the ADE2_up primer.

### Lithium acetate transformation of pML104-gRNA into BY4741 and W303

The wildtype strain of *S. cerevisiae* used in the mutagenesis experiment in this study was BY4741 while HDR was investigated in the W303-1A *ade2* mutant strain (see supplementary Material Table S2). Cells of both yeast strains were inoculated in ¼Yeast Extract Peptone Dextrose (¼YPD) broth and incubated overnight. Two 700µl aliquots of BY4741 and three 700µl aliquots of W303-1A culture were pelleted by centrifuging at 13,000rpm, the supernatants discarded, and the pellets resuspended in 355µl of transformation mixture (30% PEG4000, 100mM Lithium Acetate, 100mM 2-mercaptoethanol). The complete composition of the transformation mixture is detailed in Table 1 for the two strains.

**Table 1:**
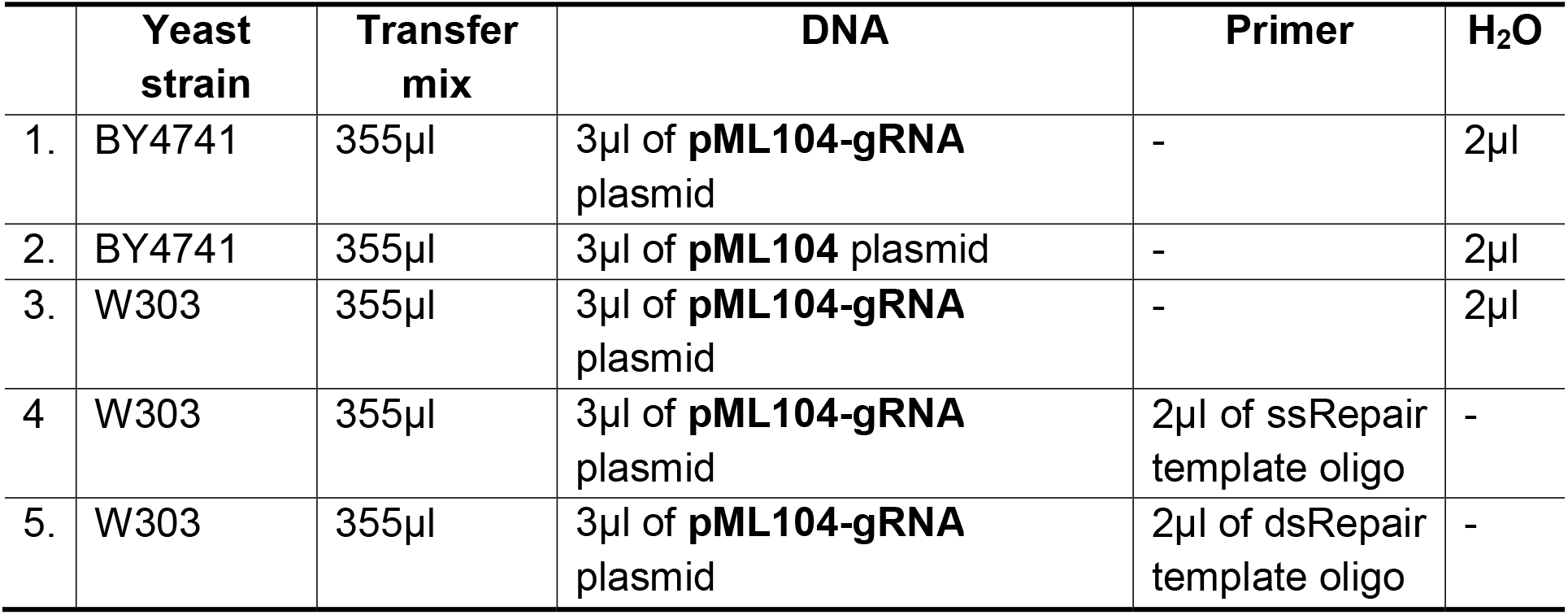
Breakdown of the four different yeast cell transformations and their constitutions. ss and dsRepair template is single- and double strand repair template oligo, respectively. The concentration of single and double stranded oligo used was 30µg/ml.ed

|  | **Yeast strain** | **Transfer mix** | **DNA** | **Primer** | **H<sub>2</sub>O** |
| --- | --- | --- | --- | --- | --- |
| 1. | BY4741 | 355µl | 3µl of **pML104-gRNA** plasmid | - | 2µl |
| 2. | BY4741 | 355µl | 3µl of **pML104** plasmid | - | 2µl |
| 3. | W303 | 355µl | 3µl of **pML104-gRNA** plasmid | - | 2µl |
| 4. | W303 | 355µl | 3µl of **pML104-gRNA** plasmid | 2µl of ssRepair template oligo | - |
| 5. | W303 | 355µl | 3µl of **pML104-gRNA** plasmid | 2µl of dsRepair template oligo | - |

To assess the efficiency of CRISPR/Cas9 mutagenesis of the *ADE2* gene, 3µl of pML104-gRNA and 3µl control plasmid (as a positive control) were separately transformed into the BY4741 cells. W303-1A cells were co-transformed with 3µl of pML104-gRNA and 2µl of either single or double stranded repair template oligonucleotides (repAde2; see supplementary Material Table S1) [Eurofins Genomics]. A negative control consisted of W303-1A cells transformed with only 3µL of pML104-gRNA. These transformation mixtures were incubated at room temperature for 20 minutes, followed by heat shock at 42ºC for 20 minutes. Cells were pelleted at 2,000rpm for 5 minutes, the supernatants were removed, and pellets were resuspended in 200µL sterile water. Each of the five yeast transformation products were then directly plated on synthetic complete (SC) uracil drop-out agar (SC–Ura) [2% glucose, 0.67% yeast nitrogen base, 0.2% Kaiser Dropout mixture without uracil (Formedium) and 2% agar]. The plates were incubated for 2 days at 30ºC.

### Verification of genome editing events

Six independent yeast transformants were randomly picked from each of the SC–Ura plates of transformations 1, 4 and 5 (Table 1). These were maintained in ¼YPD broth in separate wells of a 96-well plate. To test for the auxotrophic adenine marker of the *ade2* allele, a 5µl drop of colony-culture of each strain from these transformations, alongside reference yeast samples, were spotted onto ¼YPD agar plate (2% glucose, 0.25% yeast extract, 1% Peptone and 2% agar in water) following the arrangement shown in Figure 2.

**Figure 2:**
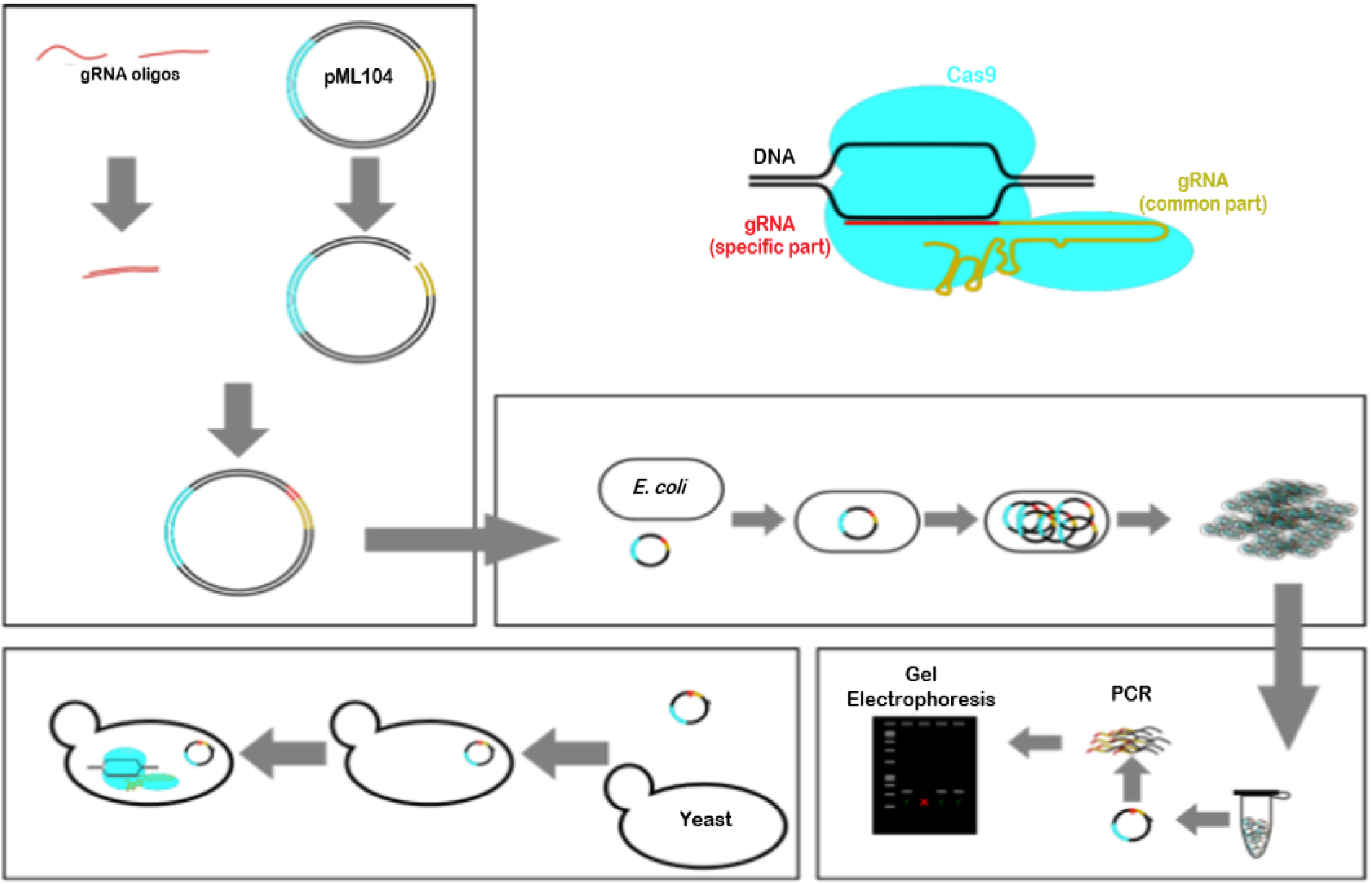
A schematic overview of the experiment. Detailed materials and methods are provided within the main text and in the Supplemental Text.

A similar spotting pattern was used for a growth assay, except 50µl of all cultures and reference yeast cells were first centrifuged for 1 minute at 13,3000rpm, and resuspended in 1ml of sterile water. The washed cells were spotted on a SC Adenine drop-out (SC–Ade) agar plate (2% glucose, 0.67% yeast nitrogen base, 0.2% Kaiser Dropout mixture without Adenine [Formedium] and 2% agar). The plates were allowed to absorb the liquid and then incubated at 30ºC for 48 hours.

## Results

### PCR confirmation of gRNA ligation into digested pML104 and bacterial transformation

The digestion of pML104 with *Swa*I and *Bcl*I resulted in linearized plasmid strands with blunt overhangs, allowing for the ligation of the annealed gRNA oligonucleotides containing compatible overhangs. Following the ligation reaction, transformation into competent *E. coli* cells resulted in colonies on LB agar plates. The number of transformants obtained from the pML104-gRNA ligation mixture was fewer than 100 (Supplementary Material Figure 1A, right), whereas the control plasmid yielded over 1000 colonies (Supplementary Material Figure 1A, left), consistent with expectations from a ligation reaction. The presence of *E. coli* transformants provided an initial indication of successful gRNA ligation into the digested pML104 plasmid. However, transformants of competent cells can still grow on a medium even if the ligation reaction was unsuccessful due to different reasons.

To confirm the insertion of the gRNA sequence, a PCR amplification of a region of the pML104-gRNA plasmid that includes the cloned gRNA was performed using primers gADE2_f (complementary to the gRNA) and g104_check (complementary to a 150bp region upstream of the gRNA sequence). The resulting PCR products exhibited an expected band size of ∼500bp, confirming the presence of the gRNA insert in the four randomly selected transformants (Supplementary Material Figure 1B). To further verify the correct orientation and position of the gRNA insert within pML104, Sanger sequencing of the clone site in pML104-gRNA plasmid was performed, followed by multiple sequence alignment against the empty pML104 using ClustalW [26]. The analysis revealed 18 nucleotide insertions and additional nucleotide substitutions in the pML104-gRNA plasmid, confirming successful incorporation of the gRNA sequence (Figure 4).

**Figure 3.**
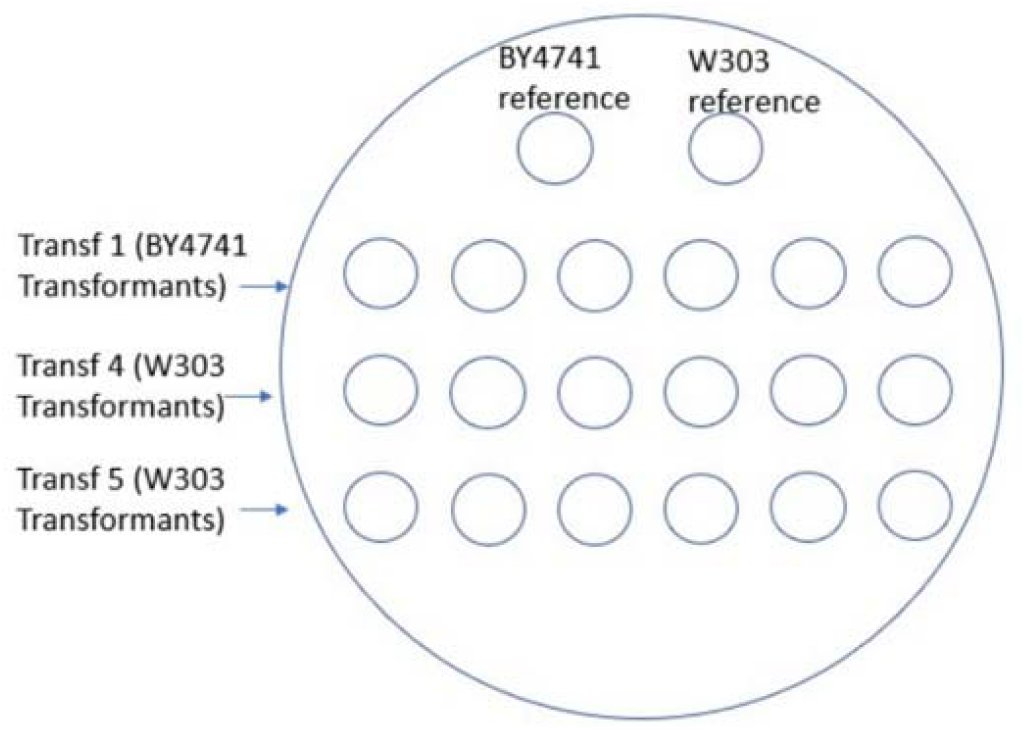
Spotting pattern for yeast colony color and growth verification assays. Reference strains represent wild-type and mutant S. cerevisiae strains that were not exposed to CRISPR mutagenesis or repair, while transformants correspond to CRISPR-modified colonies selected from SC–Ura plates.

**Figure 4.**
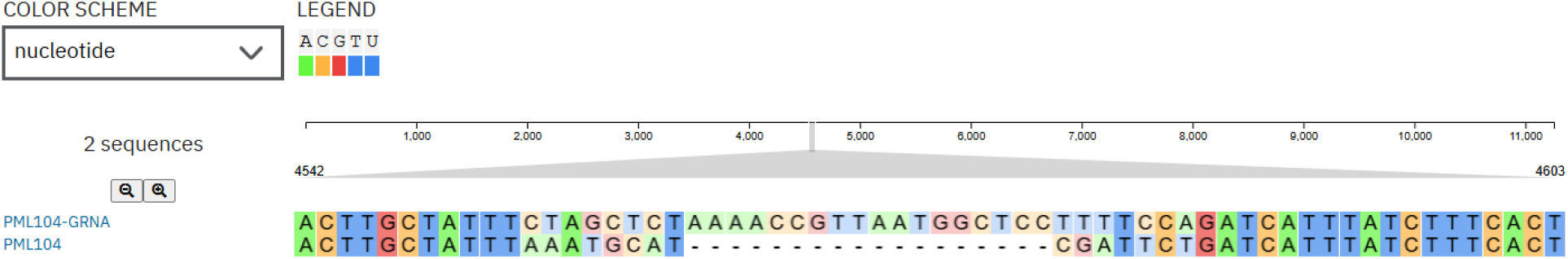
Multiple sequence alignment of the empty pML104 and the cloned pML104-gRNA using ClustalW. The observed insertions and single nucleotide polymorphisms confirm successful gRNA incorporation.

### BY4741 and W303-1A transformants

Yeast cells successfully transformed with the pML104-gRNA were selected on the SC– Ura plates, as the plasmid carries the URA3 (a yeast pyrimidine biosynthesis gene) nutritional marker. The BY4741 strain was transformed with either pML104-gRNA (transformation 1) or pML104 (transformation 2) (Table 1). The control plasmid served as a positive control to evaluate the transformation efficiency. As expected, due to the continuous gRNA-mediated cleavage and repair activities via NHEJ, the pML104-gRNA yielded fewer transformants (<50) (Figure 4A) compared to the pML104 transformants (>500) (Figure 4B). W303-1A strains require exogenous adenine otherwise their colonies accumulate the red pigment AIR. Transformation of W303-1A cells with pML104 (transformation 3) resulted in red colonies, indicating the the *ade2* mutation (Figure 4C). However, co-transformation with single- or double stranded repair oligonucleotides (transformations 4 and 5, respectively) resulted in smaller, less pigmented colonies on SC–Ura media, indicating successful HDR (Figure 4D and E, respectively).

### CRISPR/Cas9 driven mutagenesis in BY4741 and homologous recombination in W303-1A

To validate the successful CRISPR/Cas9-mediated mutagenesis of the *ADE2* gene in wildtype BY4741, a colour-based spot assay was conducted on ¼YPD medium to assess *ade2* auxotrophy. Wildtype BY4741 colonies appeared white, while transformants carrying a mutated gene exhibited red colonies, indicating successful genome editing. The mutagenesis efficiency of *ADE2* gene in BY4741 using the pML104-gRNA, calculated using the Wald correction, was 0.4 ± 0.30 (Figure 5A, Transf 1). Some BY4741 transformants remained white, suggesting either resistance to CRISPR/Cas9 editing or off-target mutations. In W303-1A, co-transformation of the pML104-gRNA with donor repair templates generated white colonies, indicative of successful HDR-mediate repair of the *ade2* gene. HDR efficiency in this study, estimated using the Wald correction, was also 0.4 ± 0.30 for both single-stranded (Figure 6A, Transf 4) and double-stranded repair oligonucleotides (Figure 6A, Transf 5). No significant difference was observed in HDR efficiency between the two repair templates; however, the statistical power was limited due to the small sample size.

**Figure 4:**
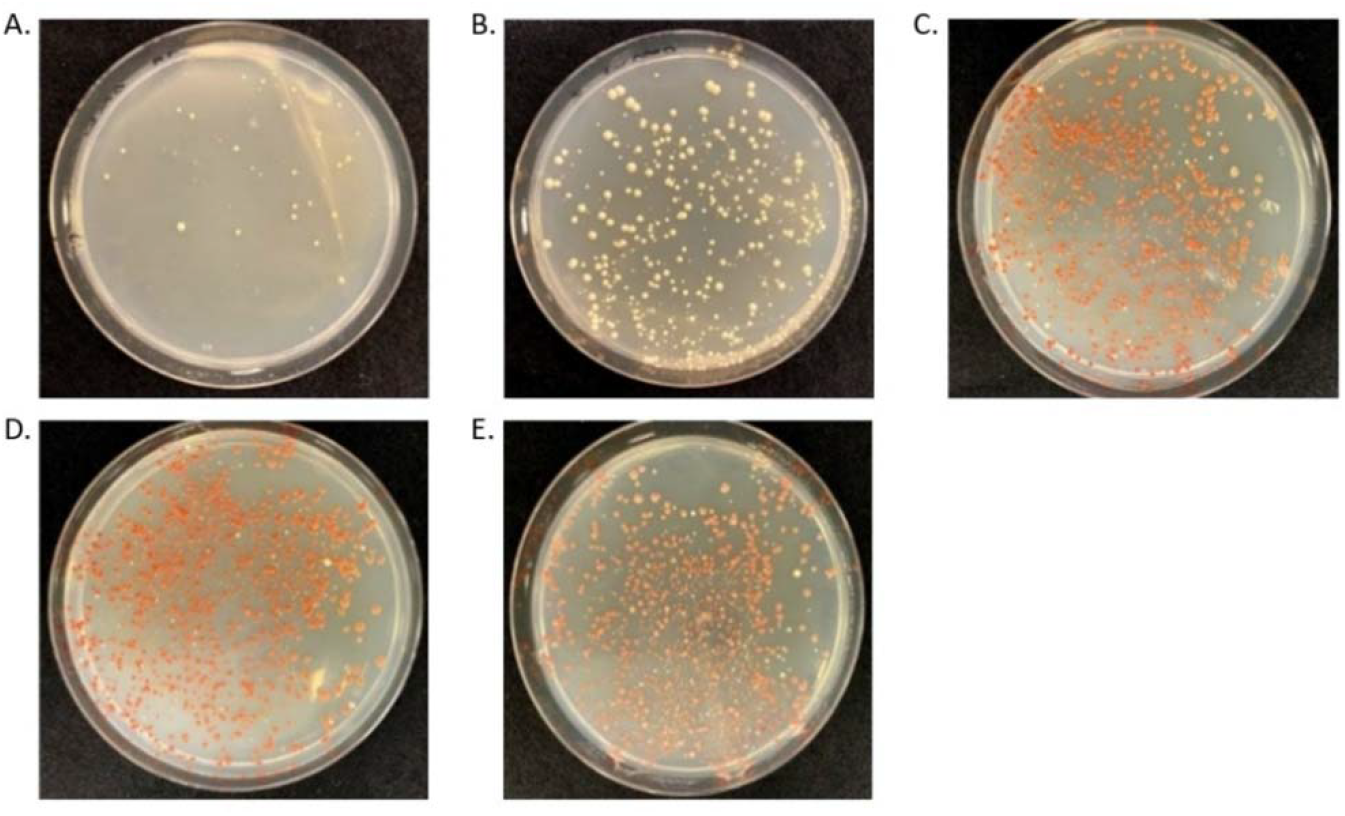
Representative yeast transformation plates corresponding to Table 1. Colony yield of BY4741 transformants carrying pML104-gRNA (A) is compared to BY4741 transformants carrying pML104 (B), demonstrating transformation efficiency. The visible red pigment accumulating in colonies of W303-1A transformed with pML104-gRNA without donor repair DNA (C), co-transformed with single-stranded repair template (D) or double stranded repair template (E). Note the changes in colony size and pigmentation with the addition of donor DNA, showing reduced pigmentation due to HDR-mediated repair.

**Figure 6.**
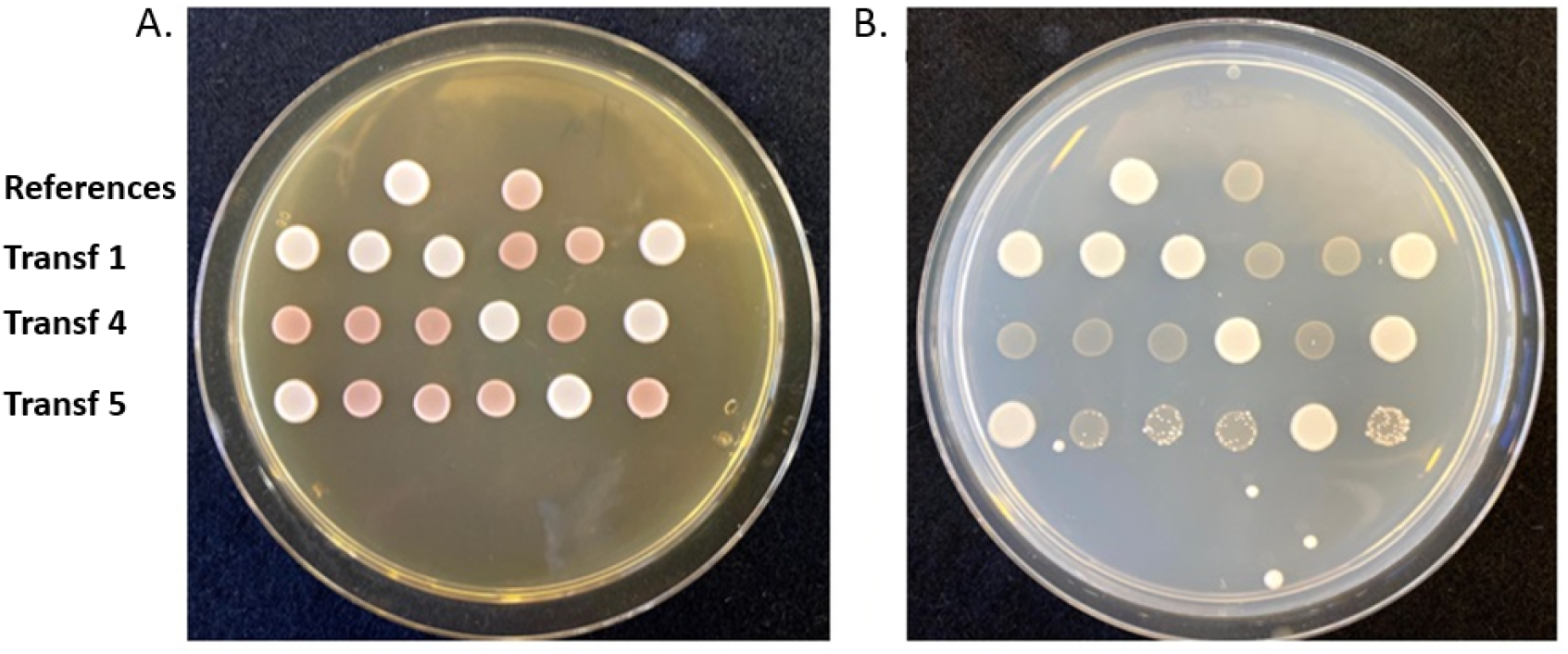
Verification of genome editing events. (A) Color assay: red BY4741 colonies confirm ade2 mutation, while white W303 colonies confirm HDR-mediated repair. (B) Growth assay: loss of ADE2 function prevents BY4741 mutants from growing on SC– Ade, while repaired W303 colonies regain growth.

To confirm that the *ADE*2 mutations in BY4741 transformants that appeared red on the ¼ YPD plate was site-specific and not off-target activities on other loci, they were plated on SC-Ade medium. Wildtype BY4741 cells were viable on the SC–Ade agar plate. CRISPR/Cas9 activity had rendered some wildtype BY4741 cells to loss their *ADE*2 gene function (corresponding to red colonies on ¼YPD) and hence failed to grow without supplementary adenine in the -Ade medium (Figure 6B, Transf 1), confirming loss of *ADE2* function. Conversely, W303-1A transformants with repaired *ade2* locus using the donor repair templates regained wildtype gene function, crucial for the activation of its adenine biosynthesis pathway on a SC–Ade agar, allowing their growth on the -Ade medium (Figure 6B, Transf 4 and 5), confirming precise HDR at the target site.

To further validate the CRISPR/Cas9 genome editing of the *ADE2* gene of the BY4741 strain, nine separate red transformants were randomly selected and prepared for Sanger sequencing to detect nucleotide deletion/substitutions. Multiple sequence alignment revealed numerous nucleotide deletions/substitutions at the *ADE2* locus in all nine BY4741 transformants (100%), characteristic of NHEJ repair (Figure 7). These changes in nucleotides rendered the *ADE*2 gene in the BY4741 strain ineffective in transcribing the accurate enzyme from the gene.

**Figure 7.**
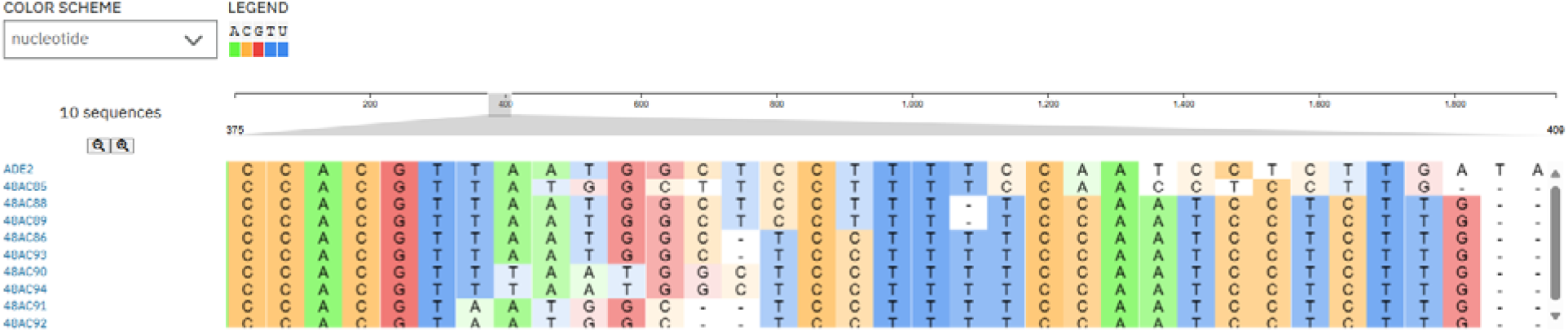
Multiple sequence alignment of the BY4741 transformants (red colonies on ¼YPD plate) generated from the CRISPR mutagenesis of the ADE2 gene in ClustalW showing deletion presentative of NHEJ repair in the transformants. Point mutation and insertions at the gRNA target site are seen.

## Discussion

CRISPR/Cas9 has vast potential as a genome editing technique in *S. cerevisiae* due to the precise targeting activity of the Cas9 nuclease, guided by user-defined gRNA sequences [27]. This study, alongside previous research, underscores the critical role of directional cloning of the 20-nucleotide guide sequence, which is complementary to the target locus, into restriction enzyme-generated breaks within the pML104 plasmid [10]. Here, Cas9 was directed to the *ADE2* locus of BY4741 by the gRNA, where it induced a DSB, leading to NHEJ-mediated repair. This process introduced sufficient mutations in BY4741, effectively disrupting *ADE2* function in 40% of its transformants. Additionally, we confirmed HDR-mediated repair of the *ade2* mutant gene when repair template oligonucleotides were co-transformed with the pML104-gRNA into W303-1A cells, allowing precise integration at the target locus.

The gRNA oligonucleotides were annealed and cloned into *Swa*I- and *Bcl*I-digested pML104, which contains the Cas9 endonuclease. The resulting pML104-gRNA functioned as an independent genome editing construct, introducing site-specific DSBs. The restriction enzyme sites in the pML104 simplified the process by eliminating the need for complex multi-step cloning of a separate gRNA and Cas9-expressing plasmid, as reported in earlier studies [27]. This approach also circumvented the necessity of assembling multiple components, such as donor DNA and gRNA sequences, into a single plasmid before transformation, as described by Bao et al. [28]. The gRNA sequence efficiently directed Cas9 to the *ADE2* locus in BY4741, where it induced DSBs that were repaired through error-prone NHEJ. The repeated target recognition-pair-cleave cycle at this locus continued until mutations accumulated to include the PAM, preventing further binding and cleavage by the Cas9 [29]. The inherent cytotoxicity of continuous cleavage-repair cycles explains the significantly lower colony count of BY4741 cells transformed with the gRNA plasmid compared to those transformed with the control plasmid. The reduced colony yield highlighted the efficiency of CRISPR/Cas9-mediated *ADE2* editing. However, some transformants retained viability on SC–Ura plates despite not acquiring the desired genomic alterations, likely due to mutations occurring outside the target locus (DiCarlo et al., 2013) or off-target effects [29].

Although NHEJ frequently introduces random mutations, co-transformation of the pML104-gRNA with donor repair templates enabled precise HDR in W303-1A cells, as documented in prior studies [10, 27]. CRISPR/Cas9-directed cleavage at the *ade2* locus continued until recombination of the repair template was successfully [29]. Interestingly, our findings indicate that single-stranded and double-stranded oligonucleotides were incorporated into the *ade2* locus with equal efficiency, contrasting with observations by Roth et al. [30].

Phenotypic differentiation of *S. cerevisiae* adenine auxotrophs via red-white colony screening [31] provides a cost-effective and rapid alternative to traditional selectable markers. The colour spot assay efficiently identified CRISPR/Cas9-edited yeast cells either through mutagenesis or homologous repair. On the ¼YPD medium, adenine depletion activates the *de novo* purine biosynthesis pathway, leading to the accumulation of AIR, which caused *ade2* mutant colonies to appear red [22, 32]. The SC–Ade medium growth assay provided an additional verification step for *ade2* mutants, as viability depends on the ability to synthesize adenine using an intact *ADE2* gene [33]. Adenine-starved yeast cells lacking functional *ADE2* undergo cell cycle arrest [22], consistent with our findings that all ade2 mutants failed to grow on SC–Ade plates.

This study successfully demonstrated CRISPR/Cas9-mediated genome editing of *ADE2* in BY4741, as well as HDR-mediated repair of *ade2* in W303-1A using a donor template. The specificity of genome editing was confirmed by Sanger sequencing, which revealed expected mutations in edited BY4741 cells. However, further optimization is needed to achieve higher rates of HDR-mediated recombination, as reported in previous studies [27, 28]. Future work should focus on improving transformation efficiencies and minimizing off-target effects to facilitate broader applications of CRISPR/Cas9 genome editing in S. cerevisiae. Despite the high predicted efficiency of the gRNA sequence using online tools, its actual performance in guiding Cas9 to introduce DSBs can be variable [29]. Increasing the concentration of the repair template during transformation or enhancing sequence homology with the target locus may improve HDR efficiency [29]. Given *S. cerevisiae* strong preference for HDR, carefully optimized experimental conditions should yield high recombination rates.

## Conclusion

Overall, our findings strengthen the utility of CRISPR/Cas9 as a powerful genome engineering tool in *S. cerevisiae*, with the methodologies outlined here potentially serving as a foundation for broader applications of CRISPR technology in microbial genetics and industrial biotechnology.

## Declaration

This study was conducted as part of the *Practical and Applied Research Skills for Advanced Biologist*s course within the MSc Infectious Diseases program at the University of Kent. As an academic requirement, it focused on applying fundamental research techniques rather than high-throughput or advanced technological approaches, and aimed to demonstrate the application of key research methodologies, and experimental design.

