## Supplementary Material for "CRISPR/Cas9-mediated editing of the *ADE2* locus in *Saccharomyces cerevisiae*: targeted mutagenesis and homology-directed repair"

Supplementary Materials

Table S1

| Oligos | Sequences |
|---|---|
| gADE2_f | GATCTGGAAAAGGAGCCATTAACGGTTTTAGAGCTAG |
| gADE2_r | TAGCTCTAAAACCGTTAATGGCTCCTTTTCCA |
| g104_check | CCGTGGTATCGTTTAGATTGGCAATTAC |
| ADE2_up | GCCGAGAATTTTGTAACACCAACATAACAC |
| ADE2_sta | CTCAATCGTTAGCACATCACATTTTTCAGC |
| repAde2 | CCAATGATCACGTTAATGGCTCCTTTTCCAATCCTCTTGATA TCGAAAACTAGCTGAAAAATGTGATGTGCTAACG |

Table S2

| Cells | Genotype |
|---|---|
| BY4741 | *MAT*a *his3Δ1 leu2Δ0 met15Δ0 ura3Δ0* |
| W303-1A | *MAT*a *leu2-3,112 trp1-1 can1-100 ura3-1 ade2-1 his3-11,15* |
| Competent *E. coli* Top10 cells | F- mcrA Δ(mrr-hsdRMS-mcrBC) φ80lacZΔM15 ΔlacX74 nupG recA1 araD139 Δ(ara-leu)7697 galE15 galK16 rpsL($Str^R$) endA1 $\lambda^-$ |

**PCR amplification cycle**

105ºC – 2 minutes - preheat lid

95ºC – 30 seconds
50ºC – 30 seconds } 30 cycles
72ºC – 30 seconds

72ºC – 5 minutes

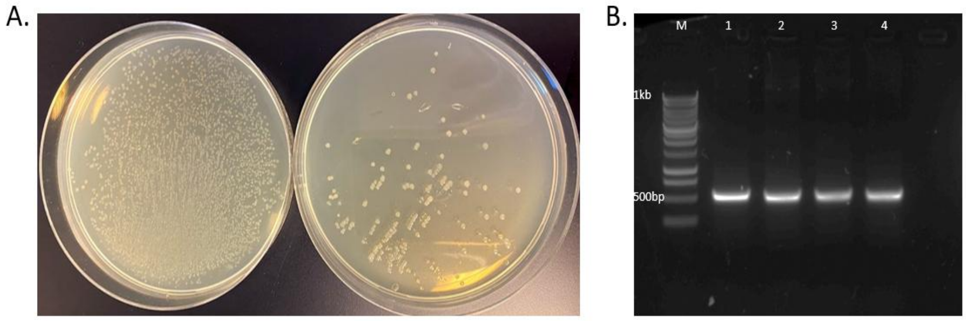


*Supplementary Figure 1: Illustrates the difference in colony numbers (A) between E. coli cells transformed with a control plasmid (left) and those transformed with the ligation mixture (right). UV transillumination of purified plasmid DNA from four randomly selected transformants*

*following PCR amplification is shown (B). A 10µL aliquot of each reaction product was electrophoresed on an agarose gel at 60V (200 mA) for 30 minutes. M represents the molecular marker (1kb DNA ladder; [Progema]), and lanes 1–4 display bands from four independent transformants.*
